# Vertical Optokinetic Eye Movements in the Larval Zebrafish

**DOI:** 10.1101/2025.03.21.644542

**Authors:** David-Samuel Burkhardt, Gabriel Möller, Laurian Deligand, Christiane Fichtner, Tim C. Hladnik, Aristides B. Arrenberg

## Abstract

The optokinetic response (OKR), a reflex enabling stable visual processing by minimizing retinal slip, has been well characterized in teleosts over the last decades. While previous work on teleost OKR mostly focused on its horizontal component, mammals are known to perform vertical and torsional OKR in addition to horizontal OKR. In this study, we characterize the vertical optokinetic response (vOKR) in larval zebrafish and compare it to the horizontal OKR (hOKR) and the vertical vestibulo-ocular reflex (vVOR). Our simultaneous camera-based tracking of vertical and horizontal eye positions reveals a distinct vOKR in larval zebrafish, but with a much smaller dynamic range compared to the hOKR and without any quick phases (resetting saccades). When presented with constant roll-rotating visual stimuli, zebrafish exhibit a brief initial vertical eye rotation in the direction of the stimulus, followed by a period without further slow phase response and interspersed with only spontaneous saccades. This behavior contrasts sharply with the periodical occurrence of resetting saccades (quick phases) during hOKR. The initial vertical response is tuned to similar spatial frequencies and angular velocities as the hOKR. We furthermore show that the vVOR has a much larger vertical dynamic range than the vOKR, demonstrating that the neuronal circuitry itself - and not the oculomotor plant - is the limiting factor. While it is unclear whether the observed differences in vertical versus horizontal optokinetic control have an adaptive value for zebrafish, the identified differences are drastic and informative for further studies on visuomotor circuits in teleosts.

**Summary Statement:** This study characterizes the vertical optokinetic response (vOKR) in larval zebrafish, revealing differences from the horizontal OKR and providing insights into visual processing and eye movement control.

## Introduction

Eye movements are ubiquitous and essential in vertebrates, enabling animals to stabilize their gaze and process visual information efficiently. Investigation of eye movements provides valuable insights into the neural circuitry underlying sensory perception, motor coordination, and behavior. In all vertebrates, eye movements are controlled by six extraocular muscles that enable horizontal, torsional, and vertical) rotations (Land, 2015), which correspond to yaw, pitch, and roll rotations, respectively, in lateral-eyed animals. It is well-established in mammals that all three directions of eye movements are used to counteract visual movements based on either vestibular or visual input, thereby stabilizing gaze position and minimizing retinal slip (Erickson and Barmack, 1980; Kitama et al., 2001; Seidman et al., 1995; van den Berg and Collewijn, 1988; Van der Steen and Collewijn, 1984).

Visual gaze stabilization is mediated via the optokinetic response (OKR) to large-field visual motion. This reflex is exhibited across vertebrate species and consists of two components: a slow phase involving relatively slow following of the visual stimulus by the eye and a resetting quick phase (a type of saccade), which moves the eye in the opposite direction of the stimulus once the maximum eye excursion is reached. The speed during the slow phase can be related to stimulus speed to calculate the gain, where unity corresponds to perfect ocular following (Dehmelt et al., 2021; Glasauer and Straka, 2022; Holm-Jensen and Peitersen, 1979; Rinner et al., 2005). The larval zebrafish is a powerful model system for studying eye movements, their underlying circuitries and development (Brysch et al., 2019; Dowell et al., 2025, 2024; Easter Jr. and Nicola, 1997; Matsuda and Kubo, 2021; Orger, 2016). This species combines a fully functional visual system, with the six extraocular eye muscles needed for movement in all directions (Bellegarda et al., 2025; Kasprick et al., 2011), and a vestibular system that enables eye movements in response to acceleration and orientation within a gravitational field (Beck et al., 2004; Lambert et al., 2008; Lim et al., 2024). Previous studies have extensively characterized horizontal (Beck et al., 2004; Dehmelt et al., 2021; Easter Jr. and Nicola, 1997; Huang and Neuhauss, 2008; Rinner et al., 2005) eye movements in larval zebrafish. Teleosts readily navigate in three dimensions in the water column, which suggests that vertical motion and vertical eye movements are important in these animals as well. Tadokoro et al. recently investigated the full repertoire of the OKR and VOR in adult goldfish (Tadokoro et al., 2025). Furthermore, vertical and torsional eye movements elicited by vestibular - not visual - stimulation have been studied in larval xenopus (Lambert et al., 2008), goldfish (Graf and Meyer, 1983; Tadokoro et al., 2025), and larval zebrafish (Bianco et al., 2012; Sugioka et al., 2023), and in archer fish 3D-motions of eyes, including vertical movements, can be measured with a stereo camera setup (Ben-Simon et al., 2009). However, visually driven vertical eye movements during the optokinetic response (vOKR), have been largely overlooked in teleost literature and only little is known about their expression and underlying circuitry.

Both optokinetic and optomotor responses stabilize fish gaze and posture based on optic flow mediated self-motion estimation (Britten, 2008; Festl et al., 2012; Kubo et al., 2014; Wang et al., 2020, 2019; Warren and Hannon, 1988). Optic flow is mainly processed in the optic tectum and pretectum by making use of a diverse set of neurons, tuned to different directions and motion patterns. Each direction-selective neuron prefers one out of four local directions: up, down, temporal-nasal (TN), and nasal-temporal (NT) motion (Gabriel et al., 2012; Hunter et al., 2013; Wang et al., 2019; Zhang et al., 2022). Although zebrafish are thought to predominantly experience horizontal motion (TN and NT) due to their swimming behavior (Ehrlich and Schoppik, 2017), the number of neurons which are direction- selective for vertical up or down visual motion is approximately equal to those encoding horizontal visual motion (Wang et al., 2019; Zhang et al., 2022). Therefore, vertical motion, be it pure up/down translation or roll or pitch rotations, seems to be very relevant for zebrafish.

In this study we simultaneously record horizontal and vertical eye movements in 4 days-old zebrafish larvae (4 days post fertilization, dpf) while eliciting both horizontal and vertical OKR by visual stimulation. Further, we quantify the vertical vestibular-ocular response, a reflex that works in concert with the OKR and relies on information from the semicircular canals and otoliths to respond to head acceleration and position. We compare the vVOR dynamics to those of the vOKR. While we show that the vOKR does exist in larval zebrafish, its characteristics strongly differ from those of the hOKR.

## Materials and Methods

### Animals

Animal experiments were permitted by the local government authorities (Regierungspraesidium Tuebingen) and conducted in accordance with German federal law and Baden-Württemberg state law. Zebrafish larvae (*Danio rerio*) were reared on a 14/10 h light/dark cycle at 28 °C in E3 medium (5 mM NaCl, 0.17 mM KCl, 0.33 mM CaCl, 0.33 mM MgSO4) with 0.01 % methylene blue. All experiments were performed on 4 dpf zebrafish larvae which are known to reliably perform hOKR (Easter Jr. and Nicola, 1997).

### Setup for VOR Experiments

To examine vertical eye movements of the vestibular ocular reflex (VOR), a setup named KEBAB (<u>K</u>inematic <u>E</u>valuation of <u>B</u>ehavior during <u>A</u>nimal <u>B</u>ody-roll) with the ability of rotating a single larva around its rostral-caudal axis (roll) was developed and built in-house. For the most part, standard Thorlabs components were used to build an examination chamber, IR-illumination for eye-tracking, visual stimulation and a camera path - all on a single axis using the 30 mm Thorlabs cage system (supplementary KEBAB). A Nema 17 stepper motor (ACT Motor GmbH, Germany) was used to rotate the whole setup. To provide electricity and enable data transfer from the steady world to the rotating camera system, its USB cable was connected via a rotating electrical slip ring (BQLZR 0.48” 12 Wire 2A 240V, Amazon). Both IR-illumination and visual stimulation consisted of LED-diodes (850 nm and white) with a 120-grid diffuser to provide uniform illumination. Recording was done at ∼20 fps using a monochromatic camera (DMK23UV024, The Imaging Source Europe GmbH). For examination, larvae were embedded on a triangle stage in 1.6 % low melting agarose as described in Wang et al. (Wang et al., 2020). Agarose around the eyes was removed to allow ocular movements (Figure S4). The stage was then clipped to a custom-made 3D-printed mount. The whole KEBAB setup was operated via an Arduino Uno with an A4988 RepRap stepper motor driver and the Python library Telemetrix (https://mryslab.github.io/telemetrix/) to enable remote control via a PC. The custom written Python-based software vxPy (https://github.com/thladnik/vxPy) was used to control setup rotation, visual stimulus presentation, camera recording and eye tracking. The noise floor of the setup was measured using anesthetized larvae (168mg/l MS222, supplementary KEBAB).

To investigate the vertical VOR, a stimulus protocol consisting of 2 x 3 stimulation phases, each framed by a 30 s pause phase was conducted in complete darkness. During the stimulation phases, the KEBAB performed 8 full rotations at either 90 °/s, 45 °/s, or 22.5 °/s velocity in either clockwise or counter-clockwise direction. Both directions were performed for each head rotation velocity. During the pause phases no rotation was performed.

### Data Analysis of VOR Experiments

First, all eye traces of stimulation phases were smoothed using a rolling mean filter with a window size of 2 s and centered around a 0 °-axis to get rid of noise and outliers. VOR score was calculated similar to Sun et al. 2018 (Sun et al., 2018) by Fourier-transforming all eye traces into the power spectrum and then taking the amplitude at the head’s rotation frequency (0.25 Hz, 0.125 Hz, 0.0625 Hz) in degree as the corresponding score. For gain and phase lag calculation, a sine-function was fitted to the stimulus triggered average (STA) of eye velocity to get the full amplitude of eye movements. Gain was then calculated as the ratio between the amplitude of eye velocity and the stimulus velocity. Phase lag was calculated by using cross-correlation.

In our vVOR experiments, we only used a single front camera whereas in our vOKR experiments two cameras (top and front) were used. During vOKR, we noticed that spontaneous horizontal eye movements occurred. Therefore, we carefully inspected the vVOR videos and confirmed that no major horizontal eye movement components occurred. This observation is mainly based on the continuous visibility of the eye lens protruding on the side in the projected front camera recording (see also Video S1).

### Setup for vOKR and hOKR Experiments

For recording horizontal and vertical optokinetic response (OKR) behavior, a modified version of the visual stimulation arena described in Dehmelt et al. (2018) (Dehmelt et al., 2018) was used. To record horizontal and vertical eye movements simultaneously, the stimulation displays in front and at the back of the fish were removed. Despite the removal of two displays, the visual stimuli of the two remaining displays still covered ∼30 % of the zebrafish field of view (Figure A6, calculated according to Dunn and Fitzgerald (Dunn and Fitzgerald, 2020)), which should suffice to elicit a reliable OKR (Dehmelt et al., 2021). An additional camera (DMK23UV024, The Imaging Source Europe GmbH) was positioned in front of the fish additionally to the camera from the top and the corresponding infrared (IR) backlight was positioned behind the fish. Fish were positioned in the middle of the arena in a water-filled translucent container and the cameras were adjusted so that both had a good view on the head and the eyes of the fish. A black lid was placed onto the container making sure that it had direct contact to the water surface to prevent unwanted reflections at the water-air surface. A similar cover was placed on the bottom of the container for the same reason. Both lid and bottom cover had a small hole in the middle to allow the IR bottom backlight to reach the fish and the camera from the top to see the fish. The two remaining displays on the left and the right side of the fish were used for OKR stimulus presentation and were sufficient to elicit OKR behavior. For each recording, a single zebrafish larva was embedded into 1.6 % agarose. The procedure was similar to that described for the vVOR experiments, but to enable both vertical and horizontal eye movements along with high-quality video recording in the IR spectrum, the agarose around and in front of the eyes was removed, leaving a small triangular portion to stabilize the head (Figure S4).

### vOKR and hOKR Stimulus Protocols

To assess whether zebrafish larvae exhibit a vOKR and characterize its tuning to different stimulus parameters, a constant rotation OKR stimulus was used. To characterize the vOKR tuning, stimulus parameters known to encompass the range of best hOKR tuning were adapted from Dehmelt et al. (2021) (Dehmelt et al., 2021). All possible combinations of six different spatial frequencies and five different angular velocities were tested (Table S1). The stimulus protocol consisted of alternating stimulation phases where the stimulus would rotate in either clockwise or counter-clockwise direction, framed by 60 s pause phases with a static grating. The order of spatial frequency and angular velocity pairs was randomized for each recording, as well as the initial direction of rotation, with each parameter pair rotating both clockwise and counter-clockwise.

Since larvae only responded within the first few seconds to a constant rotating stimulus, a second stimulus protocol using sinusoidally modulated rotation was designed with the aim of eliciting a more robust and prolonged vOKR. The number of stimulus periods within one second is defined as the repetition rate. First, vOKR tuning to 10 different repetition rates was examined, keeping the angular velocity and spatial frequency constant at 12.5 °/s and 0.0611 cycle/°, respectively (Table S2). Each repetition rate was displayed twice in a randomized order and with randomized directions, with one stimulation phase consisting of 6 stimulus periods. Stimulation phases were framed by 20 s pause phases with a static grating.

After characterizing the vOKR tuning to repetition rate, the vOKR tuning to angular velocity and spatial frequency was assessed, using the same parameter range as for the constant rotation stimulus protocol with the sinusoidal modulation at repetition rate of 0.125 Hz (Table S1).

### Data Analysis of vOKR Experiments

All eye traces were filtered with a median filter with a window size of 1 s to remove noise while preserving the velocity profile of the saccadic eye movements. Saccadic eye movements were detected using a multi-step process, including identifying the start and end points of each saccade based on the first and second derivatives of the eye traces laying within a time window of 0.5 s. A threshold of 1 ° minimal deflection and a threshold of 30 °/s or 10 °/s peak velocity during saccadic movements (for horizontal and vertical eye movements respectively) and a threshold of 20 °/s or 10 °/s average velocity (for horizontal and vertical eye movements respectively) were applied, with the values selected based on visual inspection of the data. If multiple potential start and end points were detected for a single saccade, amplitude, average velocity, change in direction of eye movements, and peak acceleration were used with different weightings to determine the most appropriate start and end points. This was the same for both recordings using the constant and sinusoidal rotating stimulus. After start and end timepoints of each saccade had been determined in a last step, the frames before and after both start and endpoint were screened for eye deflection differences. If in the frame before or after start or end a more extreme eye deflection was reached that exceeded 0.5 °, the time point for start or end was shifted to the frame with the eye trace extremum. Saccades were detected individually for each eye and camera.

The gain during the constant rotation stimulus was calculated by taking the ratio of the eye deflection with removed saccades within the first four seconds of each stimulation phase to the corresponding stimulus position. Cumulative average eye positions were calculated by removing all saccades and then averaging all eye traces across all stimulation phases. This same process was applied to the first 15 seconds, but without removing saccades, to better illustrate the initial vertical response.

Recordings performed alongside the sinusoidal rotating stimulus were often disrupted by spontaneous saccades, although showing an overall sinusoidal pattern. A combined Wiener and median filter were applied to remove high-frequency eye movements, and the signal was detrended afterwards. All further analysis was done using the average response to one stimulus period (stimulus triggered average - STA). Eye response periods where 50 % of all data points exceeded the standard deviation over all other periods within the same stimulation phase were removed from the STA to account for potential distortions by left over saccades. Gain was calculated by taking the ratio of a peak-to-peak sine function plus linear function fitting over the STA and the peak-to-peak stimulus position (similar to Dehmelt et al., 2021 (Dehmelt et al., 2021)). Since the STA of long phases (small repetition rates) exhibited a plateau of maximum eye deflection which would drastically impair the fit of the sine curve, these plateaus were excluded from the fitting process. For vOKR and hOKR (but not vVOR), we fitted the sinusoidal function to the eye position STA instead of the eye velocity STA, because eye position STA were less noisy. The fitting results were similar (data not shown).

Dynamic range was calculated as the difference between the maximum and minimum eye positions of the STA. The top of descent (ToD), used for analyzing phase lag, was defined as the point where a rolling mean of the STA gradient decreased by half of the standard deviation after the peak of the sine curve. The phase lag was then calculated as the difference between the ToD of the stimulus and the ToD of the STA.

To analyze different saccade types, all saccades during vOKR and hOKR constant rotation stimuli were sorted corresponding to their apparent direction components into four clusters (Figure S3A and B). When horizontal and vertical movements were detected on both horizontal and vertical eye trace and occurred within a time window of 5 frames, they were assigned to the same saccade and sorted in the corresponding clusters 3 or 4. If no corresponding saccade was found in the other eye trace, the saccade was characterized as only vertical or horizontal, respectively (clusters 1 or 2). In rare cases, multiple saccades occurred in one camera eye trace within the window and were therefore not assigned to any cluster. To verify the correct detection of pure vertical and pure horizontal saccades, the change of the minor axis length (i.e. the minor axis of the fitted ellipse)was calculated, taking the difference between the axis length before and after the saccade in pixels and dividing it by the major axis length of the eye to norm it (Figure S3C). The normalization step was necessary since the overall axis length is in pixels and therefore also the overall change varied between larvae. The difference images of pre and post saccade eye positions were calculated using ImageJ (Schindelin et al., 2012).

## Results

### vVOR has a large dynamic range and is driven by rotational head position

To demonstrate that zebrafish larvae are capable of large vertical eye movements, we first performed vVOR experiments. Animals were mounted in 1.6 % agarose in a setup that allowed controlled rotations about the caudo-rostral axis (clockwise and counter-clockwise roll; CW and CCW) alongside tracking of vertical eye movements. Vertical eye movements were recorded in the dark via a camera placed on the same rotation axis (Figure 1A). Three different angular velocities (22.5 °/s, 45 °/s and 90 °/s) were tested to examine vertical eye movements elicited by vestibular input. In total n = 9 larvae were used in this experiment. Throughout the experiments, the larvae exhibited a nearly perfect sinusoidal pattern of vertical eye position changes in response to the full rotations about the caudo-rostral axis, with the maximal eye deflection occurring at 90 ° and 270 ° head positions (Figure 1B and Video S1). Occasionally, the eye traces contained small outliers, which could be traced back to irregularly occurring vibrations of the stepper motor driving the setup. The average dynamic range of both eyes increased with head velocity, reaching a maximum at 90 °/s (32.3 ° ± 6.4 °; 29.3 ° ± 6.2 ° and 27.0 ° ± 5.9 ° for 45 °/s and 22.5 °/s, respectively; mean ± SD) (Figure 1C). Interestingly, the larvae did not perform any saccades during the head rotations, while the saccade rate during the pause phases, when no rotation was performed, was 0.82 saccades/min (Figure 1D). To analyze only the components of the vertical eye movements related to the stimulus frequency, Fourier analysis was performed to calculate the VOR score. The VOR scores differed with regard to the angular velocities but were nearly normally distributed for all angular velocities, with means of 15.9 ° ± 3.4 °, 13.9 ° ± 3.4 °, and 13.0 ° ± 3.3 ° (mean ± SD for 90 °/s, 45 °/s and 22.5 °/s angular velocity) (Figure 1E). The fact that the vVOR score (15.9 °, 13.9 ° and 13.0 °; mean) and the dynamic range half amplitude (16.2 °, 14.7 °, 13.5 °; mean) were close to each other indicates that the eye movements were almost solely driven by the rotational head position. The vVOR gain revealed differences across the angular velocities, peaking at 0.33 for 90 °/s (Figure 1F). The phase lag was small throughout (< 2 °), suggesting that the eyes nearly simultaneously moved together with the head, with almost no delay (Figure 1G).

**Figure 1:**
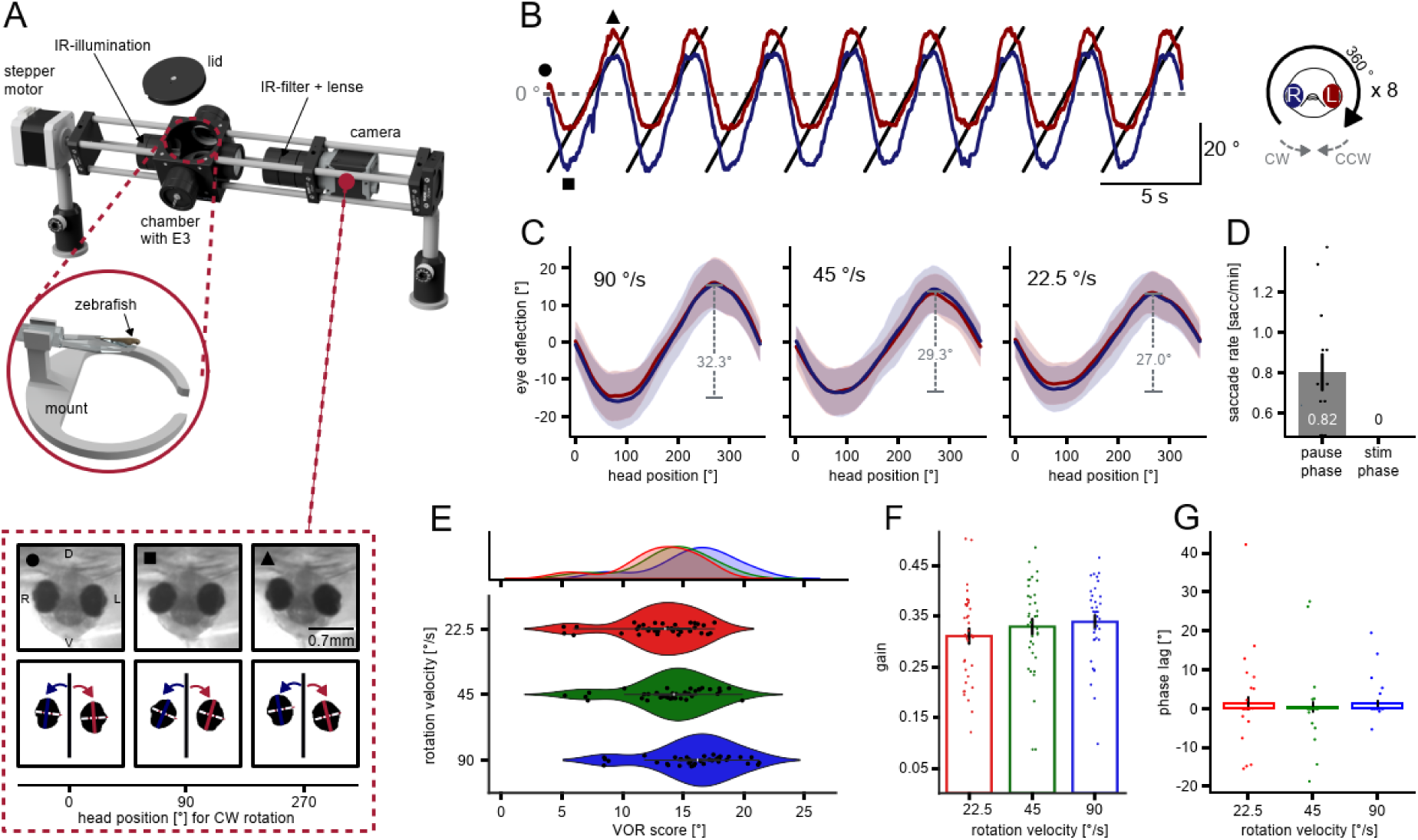
Vertical vestibulo-ocular reflex in larval zebrafish. **(A)** Schematic of the experimental setup (KEBAB) used to examine vertical eye movements during constant rotation of the fish (KEBAB is an abbreviation for <u>K</u>inematic <u>E</u>valuation of <u>B</u>ehavior during <u>A</u>nimal <u>B</u>ody Roll). The setup was driven by a stepper motor and controlled via the open-source software vxPy. Bottom: illustration of tracking vertical eye movements. **(B)** Raw eye traces of the left and right eye during rotation at 90 °/s. The symbols (dot, square and triangle) indicate the eye position displayed in panel A). One stimulation phase contained 8 full rotations, either CW or CCW. **(C)** Stimulus triggered average (STA) of eye positions during full rotation at 90 °/s, 45 °/s and 22.5 °/s for all stimulation phases and across all fish (n = 9). Envelopes show the standard deviation. Grey dashed lines indicate the dynamic range. **(D)** Saccade rate during rotation and pause phase. **(E)** VOR score for 90 °/s (blue), 45 °/s (green) and 22.5 °/s (red). VOR score only contained eye movements at the head rotation frequency. **(F)** Gain as ratio of sinus-fitted eye velocity to head velocity. **(G)** Phase lag as temporal difference between sinus-fitted eye velocity to head velocity.

### vOKR has much smaller dynamic range than hOKR and vertical quick phases are absent

To investigate the existence and functionality of the vOKR in zebrafish, a horizontally striped black and white pattern was rotated vertically around the fish at a constant angular velocity. Eye movements were tracked in two dimensions using top and front cameras to record both horizontal and vertical eye movements (Figure 2A). Various angular velocities and spatial frequencies were tested to assess the vOKR (Table S1). Each combination of stimulus parameters was presented for 60 seconds, followed by a 60-second pause phase (Figure 2B). When presented with a constantly rotating vertical optokinetic stimulus, the zebrafish larvae did not exhibit the typical pattern of slow and quick phases observed in the hOKR. Instead, the eyes followed the visual stimulus for the first 4-5 seconds, reaching a plateau-like position from which they did not move further. This “start response” was primarily visible in the frontal camera, suggesting it was limited to the vertical component (Figure 2C). After reaching the plateau, the eyes began performing spontaneous saccades in both clockwise and counter-clockwise (with and against the stimulus direction) directions, which were visible in both the frontal and top cameras, indicating these saccades might have both vertical and horizontal components (Figure 2B). Between the spontaneous saccades, the eyes remained stationary or slowly drifted back towards a more neutral position. On average, the vertical plateau-like position was held during stimulation with the eyes drifting back to neutral positions in the subsequent pause phase (Figure S1A). To visualize this surprising “start response” behavior, we compared the beginning phases of vOKR and hOKR (Figure 2D). For hOKR we used the same stimulus protocol but with vertical stripes and horizontal instead of vertical rotation (Figure 2D). When removing all saccades during vertical and horizontal OKR and calculating the average cumulative eye trace during stimulation, the hOKR revealed the well-known constant slope in the direction of the stimulus, while the vOKR flattened out after about 4-5 seconds. Vertical eye movements during the horizontal stimulus protocol were also apparent, as observed in the front camera (Figure 2D). The presence of larger eye position changes in the front camera during hOKR suggests that larval zebrafish either perform a small vertical component during the hOKR or that the projection of the eye onto the front camera is systematically affected by horizontal eye position changes. For the vOKR, horizontal eye position (measured by the top camera and removing saccades) remained stable around zero during vertical eye movements (Figure 2D).

**Figure 2:**
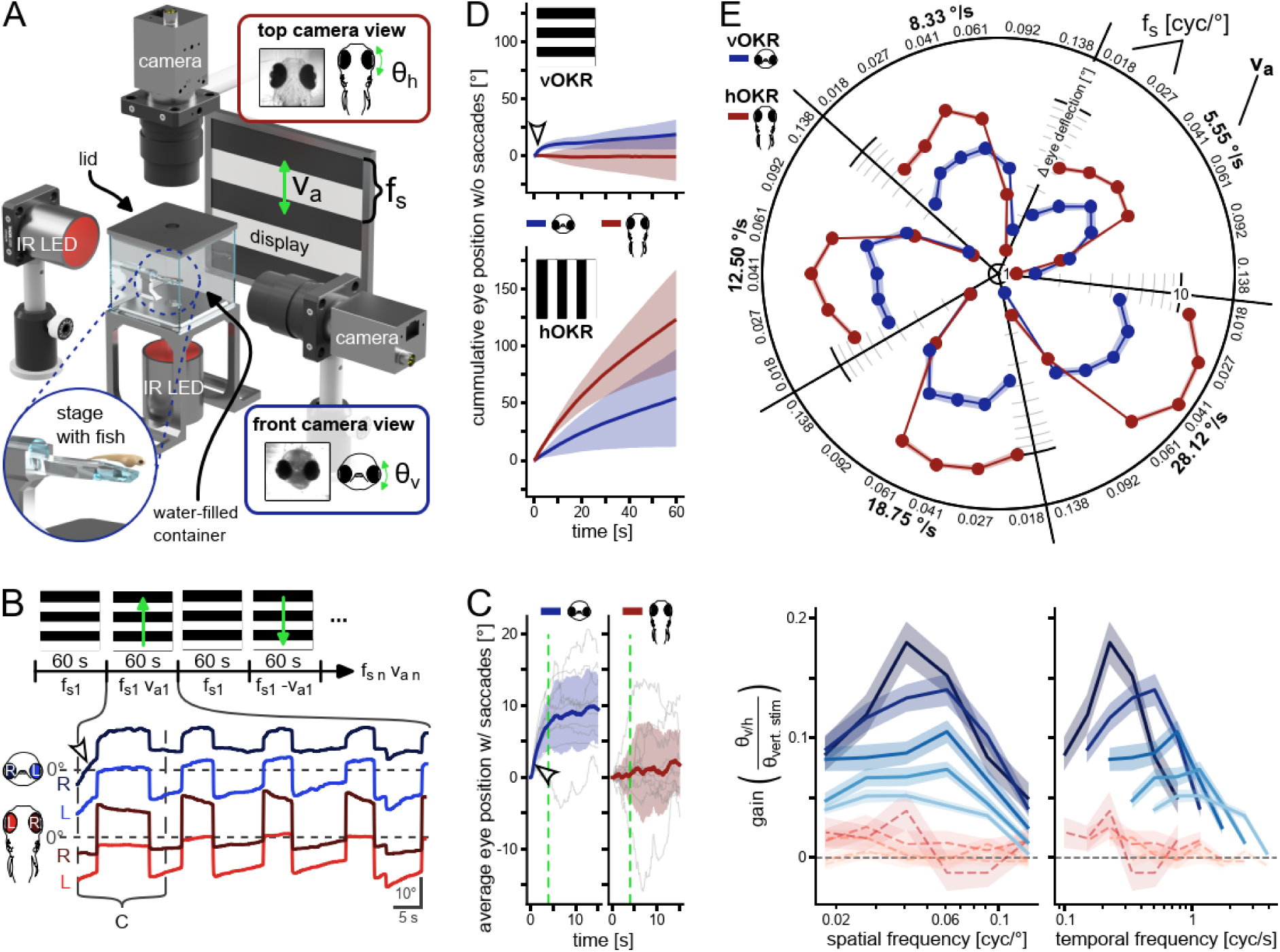
No vertical quick phases after initial zebrafish vOKR. **(A)** Schematic of the setup with two cameras simultaneously recording the horizontal (top) and vertical (front) eye movements. The second display on the right side of the fish is not shown in this illustration. **(B)** Stimulus protocol and example vertical (blue) and horizontal (red) eye traces of one stimulation phase of one fish. During the first seconds, only the vertical eye trace shows a deflection (arrowhead). Vertical dotted lines indicate cutout of panel C. **(C)** Average eye position of all fish and all visual stimulations for the first 15 s. Vertical eye trace (blue) shows a distinct rise (arrowhead) in the first 4 seconds (green dashed line) while the average horizontal eye position (red) does not change during vOKR. **(D)** Average cumulative eye position with saccades removed for vOKR (top) and hOKR (bottom) stimuli. The vertical start response is visible (arrowhead). Blue lines indicate front camera readings (vertical eye trace), red lines horizontal eye traces. **(E)** Mean amplitude within first 4 s after stimulus onset and standard error of the mean for each stimulus parameter pair. Blue: vertical amplitude for vOKR, red: horizontal amplitude for hOKR. **(F)** Gain tuning to spatial frequency and temporal frequency for all different angular velocities. Lines show mean and standard error of the mean. Blue: gain for vertical eye trace for vOKR stimulation. Red: gain calculated for horizontal eye trace for vOKR stimulation. Dashed horizontal line indicates a gain of 0. n = 10 fish.

To examine the tuning of the start response during vOKR for different spatial frequencies and angular velocities, and to compare it to the tuning of hOKR, the eye position amplitude (half the dynamic range) and the gain were calculated for the first 4 seconds of stimulation, after which the eye deflection reached the previous described plateau. The dynamic range tuning was quite similar for vOKR and hOKR (Figure 2E). For all angular velocities, the dynamic range was around zero for the highest spatial frequency of 0.138 cycles/°, then increased with lower spatial frequency until reaching a maximal deflection for either 0.061 cycles/° or 0.041 cycles/° (hOKR and vOKR respectively) before starting to decline again. Additionally, the dynamic range of eye positions increased with angular velocity, with the dynamic range being more narrowly tuned during hOKR than during vOKR. The maximum amplitude for hOKR after 4 s was 16.92 ° ± 0.61 ° (at 28.12 °/s and 0.041 cycles/°), and 5.81 ° ± 0.38 ° for vOKR (at 28.12 °/s and 0.027 cycles/°).

The vOKR gain was generally higher for slower angular velocities (Figure 2F). Within a single angular velocity, the gain showed unimodal tuning in the range from 0.018 cycles/° to 0.138 cycle/° and peaked at medium spatial frequencies around 0.04 cycles/° to 0.06 cycles/°. The same pattern held true when comparing the gain to temporal frequencies, where the slower the angular velocity, the lower the temporal frequency that elicited the highest gain. To further confirm that the eye movements recorded within the first seconds of stimulation only contained a vertical OKR, the gain was also calculated for the horizontal eye movements during vOKR stimulation. The resulting values all lay around zero, indicating no notable horizontal component was present in the early part of the vOKR.

### Periodic vOKR shows phase shifts depending on the time period to stimulus reversal

Since the constant roll-rotating visual stimulus only elicited a transient vOKR within 4-5 seconds, a second sinusoidal rotation stimulus was used to extend the duration for which the eyes would follow the visual motion. The stimulus contained the same striped black and white pattern, but the rotation velocity followed a sinusoidal profile (Figure 3A). As a result, the stimulus slowly gained speed in one direction until reaching the maximum angular velocity, then decreased again until coming to a full stop before accelerating in the opposite direction. This stimulus introduced a new parameter, the repetition rate (rr), which indicated the frequency of the sinusoidal velocity envelope and provided information on how quickly the maximum velocity was reached. Overall, the sinusoidal stimulus elicited sinusoidal vertical eye movements corresponding to the stimulus position, while no stimulus-associated horizontal eye movements were visible in the top camera (Video S2). During stimulation, larvae showed spontaneous horizontal saccades that also led to deflections in the vertical eye position but did not disrupt the overall sinusoidal pattern. Just as for the constantly rotating stimulus, no vertical resetting quick phases could be observed in response to the sinusoidal stimulus (Figure 3B).

**Figure 3:**
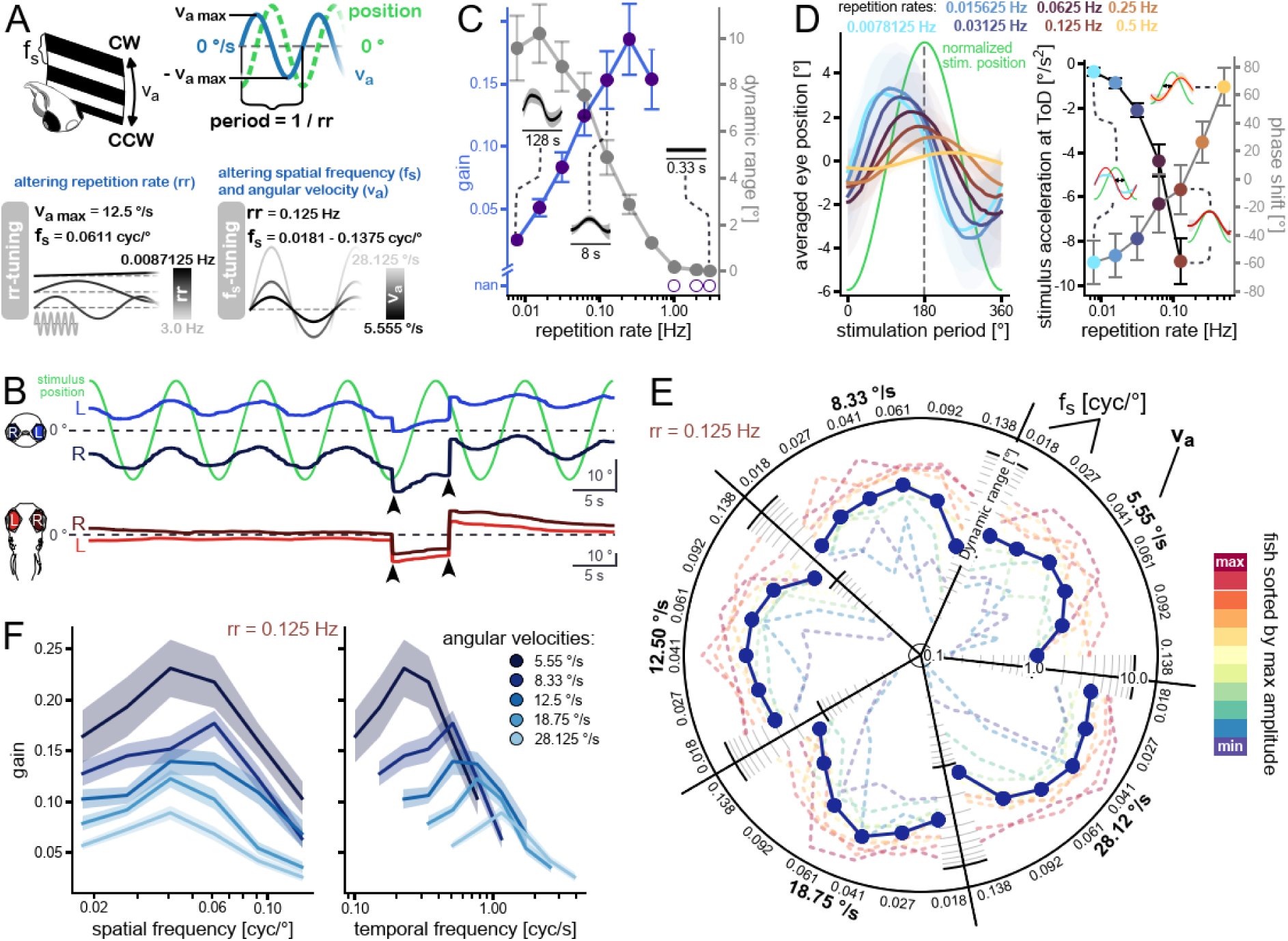
vOKR tuning to stimuli with sinusoidal velocity envelope. **(A)** Sinusoidal rotation stimulus scheme. Two different sinusoidal stimuli were used. In the rr tuning stimulus, the maximum angular velocity and the spatial frequency were constant while different repetition rates were tested. In the f_s_ tuning stimulus, the repetition rate was kept constant while spatial frequency and angular velocity were altered. CW: clockwise, CCW: counterclockwise, v_a_: angular velocity, rr: repetition rate, f_s_: spatial frequency. **(B)** Example recording with vertical (blue) and horizontal (red) eye traces. Spontaneous saccades during the sinusoidal response did not disturb the response but shifted it by the amplitude of the saccade (arrowheads). Note: different y-axis scales. **(C)** Mean and standard error of gain (blue) and full dynamic range (grey) over repetition rate. Insets show average response to one stimulus period over all fish for repetition rates 3.0 Hz, 0.125 Hz, and 0.0078125 Hz (n = 8 fish). **(D)** Left: Average response to one stimulus period for different repetition rates. The timescale is normalized to one stimulus period. Vertical dashed line indicates turning point of stimulus position. Right: Phase lag of the vOKR (gray lines) for different repetition rates; and acceleration value of the stimulus (black lines) at the time point of eye movement direction change (phase lag). Data Points are color-coded with respect to their repetition rate and show mean and standard deviation. Insets show example average responses to one stimulus period of one fish. Green curve indicates stimulus position, red curve shows sine fit to average eye trace. Black arrows indicate the phase lag between stimulus and average response **(E)** Dynamic range for each stimulus parameter set. Datapoints indicate mean of all fish (n = 11 fish). Dashed lines show mean dynamic range of single fish, colored according to their mean amplitude. **(F)** Mean gain with standard error over spatial frequency (left) and temporal frequency (right) for all different angular velocities (see color legend).

We tested different repetition rates while keeping the maximum angular velocity constant (12.5 °/s) and using only one spatial frequency close to the optimum (0.061 cycle/°) (Table S2). The parameters were similar to those used by Dehmelt et al. (2021) for horizontal OKR (Dehmelt et al., 2021). For repetition rates between 0.5 Hz and 0.0078 Hz, vertical eye positions followed the stimulus. Repetition rates greater than 0.5 Hz did not elicit any response (Figure 3C and S2A). The gain was calculated as the ratio of the amplitude of a fitted sinusoidal function to the eye position and the amplitude of the sinusoidal function of stimulus position. The gain was relatively high for repetition rates between 0.5 Hz and 0.125 Hz (maximum of 0.19 at 0.25 Hz), but started to decrease for lower repetition rates until reaching a value of 0.03 (at 0.0078 Hz) (Figure 3C). However, the eye traces themselves showed a clear sinusoidal pattern for low repetition rates, and the dynamic range even increased with lower repetition rates. For the highest repetition rates that did not elicit any response, the dynamic range was around zero, but reached around 10° for the lowest two repetition rates. Analyzing the stimulus triggered average (STA) of eye position for all repetition rates revealed a phase lag with regard to stimulus position, which moved from positive phase angles towards negative ones with decreasing repetition rates (Figure 3D). For high repetition rates (0.25 and 0.5 Hz), the eyes therefore seemed to lag behind the stimulus, which is expected. For lower repetition rates (0.0078 to 0.125 Hz), however, the eye movement preceded stimulus position change, which is counterintuitive. This negative phase lag meant that the eyes already started moving in the opposite direction during the slow-down of the current stimulus (before the sign of the stimulus velocity would change). For the highest repetition rate with responses (0.5 Hz), the phase lag was 65.76 ° ± 13.31 °, while for the lowest repetition rate (0.0078 Hz), the phase lag was -59.37 ° ± 15.69 ° (i.e. a phase advance). When analyzing those phase advances with regard to the eye’s acceleration, we found that the sign of eye velocity only changed during slow-down of the stimulus, i.e. after the acceleration turned negative (Figure 2D, right panel). This phenomenon can potentially be explained by the motion aftereffect or acceleration coding neurons and will be discussed further below.

The vOKR to a repetition rate of 0.125 Hz had the smallest phase lag and was used to investigate the tuning to spatial frequency. The angular velocities and spatial frequencies were varied following the constant rotation stimulus protocol (Table S1). The dynamic range of vertical eye movements during sinusoidal vOKR overall revealed a similar pattern compared to the constant rotating stimulus (Figure 3E). On average, the dynamic range of eye position was small for high spatial frequencies, then began increasing and after peaking at 0.041 cycle/° or 0.061 cycle/°, it declined again for lower spatial frequencies. The maximum dynamic range for the slowest angular velocity (5.55 °/s) was 3.31 ° ± 2.44 °, and the maximum dynamic range for the fastest angular velocity (28.12 °/s) was 6.21 ° ± 3.02 °. When examining individual fish, much higher dynamic ranges, up to >10 ° (red dashed lines, Figure 3E) occurred in some fish. Two fish exhibited very distinct dynamic range tuning properties that stood out from the rest (purple and blue dashed lines). For high spatial frequencies, their dynamic range was much smaller (<1 °) than in the other fish. Furthermore, their maximum dynamic range for slower angular velocities was also relatively small, only starting to match the others as angular velocities increased.

The tuning of vOKR to spatial frequency and temporal frequency was similar for sinusoidal and constant rotation stimuli. The overall gains were higher for the sinusoidal rotation stimulus, though, reaching a maximum gain of 0.23 ± 0.03 (at 5.55 °/s and 0.041 cycle/°, see Figure 3F).

### Zebrafish larvae generate pure vertical and pure horizontal spontaneous saccades

In our recordings, saccadic eye movements were detected during both vOKR and hOKR stimulation. During vVOR, no saccades were visible. Saccadic movements occurred in both the horizontal and vertical planes. Interestingly, the saccade rate during vOKR stimulation was similar to the rate during no stimulation, suggesting these were purely spontaneous (Figure 4A). Furthermore, neither the stimulus direction nor the type of stimulation (constant vs. sinusoidal) influenced the saccade rate during vOKR stimulation. During hOKR, most saccades occurred against the direction of the stimulus, mirroring the well-known pattern of resetting quick phases during stimulation. To further characterize saccadic eye movements during vOKR stimulation, saccades were sorted into four clusters based on their occurrence with or without corresponding eye movements in the other plane (Figure S3A and B). A total of 4696 detected saccades were analyzed during vOKR. The first cluster contained saccades with only a vertical component (10.5% of all saccades), while saccades with only a horizontal component were sorted into cluster two (14.0% of all saccades). Saccades with apparent deflections in both the vertical and horizontal planes were sorted into clusters three and four, depending on their direction (75.5%). Large deflections in the horizontal plane caused a more spherical (less elliptical) appearance of the eye in the frontal camera. This projection artifact often effected false estimates of vertical eye position, since the detected major axis angle was affected by the ellipse fit to the now more spherical shape. Therefore, it was virtually impossible to extract the true vertical and horizontal components of apparently combined eye movements. For this reason, clusters three and four were not further analyzed, and they likely contain pure horizontal saccades and/or true vertical-horizontal combined eye movements (but no pure vertical saccades).

**Figure 4:**
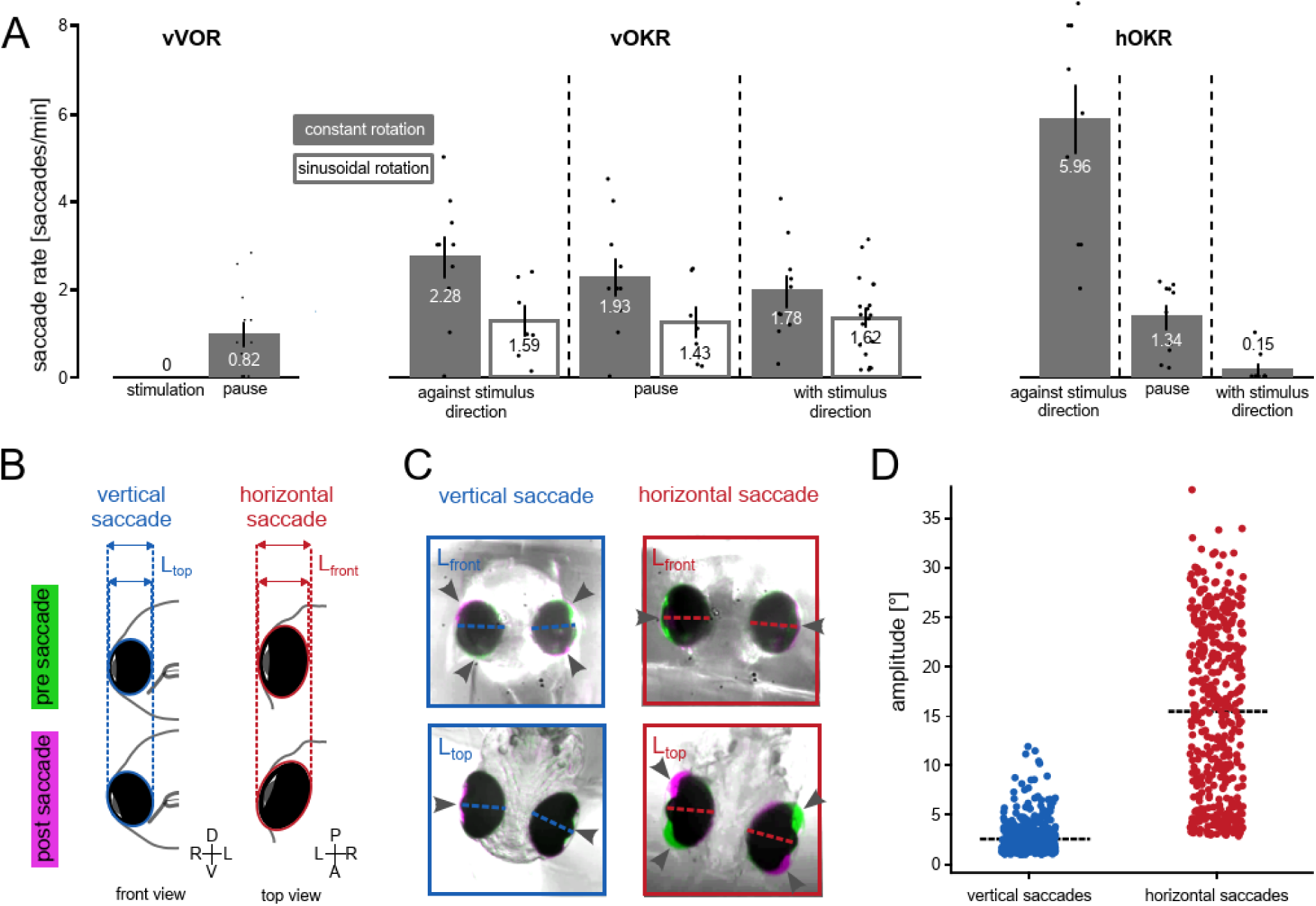
Characterization of vertical and horizontal saccades. **(A)** Comparison of the saccade rate (vertical + horizontal saccades) during vVOR, vOKR and hOKR. For all stimuli, data from the stimulation parameters that elicited the highest number of saccades are shown **(B)** Schematic of minor axis length change (ellipse fitting) during vertical and horizontal saccades. During vertical saccades, the minor axis length in the top camera is expected to change, whereas the minor axis length in the front camera should remain constant. For horizontal saccades the expectation is vice versa. These length changes together with the angle between the major axis and the body axis were used to detect pure vertical and horizontal eye movements (see also Figure S3). **(C)** Difference image of the eyes’ pre-saccadic (green) and post-saccadic (magenta) positions for pure vertical and pure horizontal example saccades. Arrows indicate the area where position changed. For pure vertical saccades, the top camera showed changes only at the lateral parts of the eyes, whereas the front camera exhibited changes at the dorsal and ventral parts of the eyes. Conversely, during pure horizontal saccades, the image changes were reversed (change at lateral parts of eyes in front camera and at anterior/posterior parts of eyes in the top view camera). **(D)** Saccade amplitude scatter plot of pure vertical and pure horizontal saccades.

To ensure that clusters 1 and 2 exclusively contained pure vertical (Video S3) and pure horizontal (Video S4) saccades, and that saccadic eye movements within a single plane were not affected by the above mentioned projection artifact, the change in length of the eye’s fitted minor axis was examined (Figure 4B). For pure vertical movements, the minor axis length was expected to change in the horizontal plane (top view) and stay constant in the vertical plane (front view). Conversely, during pure horizontal eye movements, the minor axis length change was expected to occur in the vertical plane while remaining at zero in the horizontal plane (Figure SD). Analysis of the minor axis change distribution during both pure vertical and pure horizontal saccades yielded the expected results, thereby confirming the accurate detection of pure vertical and pure horizontal saccades. To illustrate examples of pure vertical and horizontal eye movements, false color images of pre- and postsaccadic camera recordings were created (Figure 4C).

Pure vertical saccades almost never exceeded 10° in magnitude but frequently exhibited a slight horizontal deflection of a few degrees (Figure S3B). In contrast, pure horizontal saccades had much larger amplitudes, reaching over 30° for some of them (Figure 4D).

### Comparison of vOKR, hOKR and vVOR performance levels

Our findings are summarized in the comparison chart in Figure 5. Comparing the vOKR and vVOR is especially intriguing, as both involve the same extraocular eye muscles but are driven by different sensory inputs. While resetting saccades are a characteristic feature of the hOKR, they did not occur during the vOKR or vVOR. Interestingly, spontaneous saccades were regularly observed in both directions (against and with stimulus direction) during the vOKR, whereas no saccades were present during the vVOR, as indicated in Figure 4A. This suggests that the vestibular roll stimulation or vVOR suppresses spontaneous saccadic eye movements.

**Figure 5:**
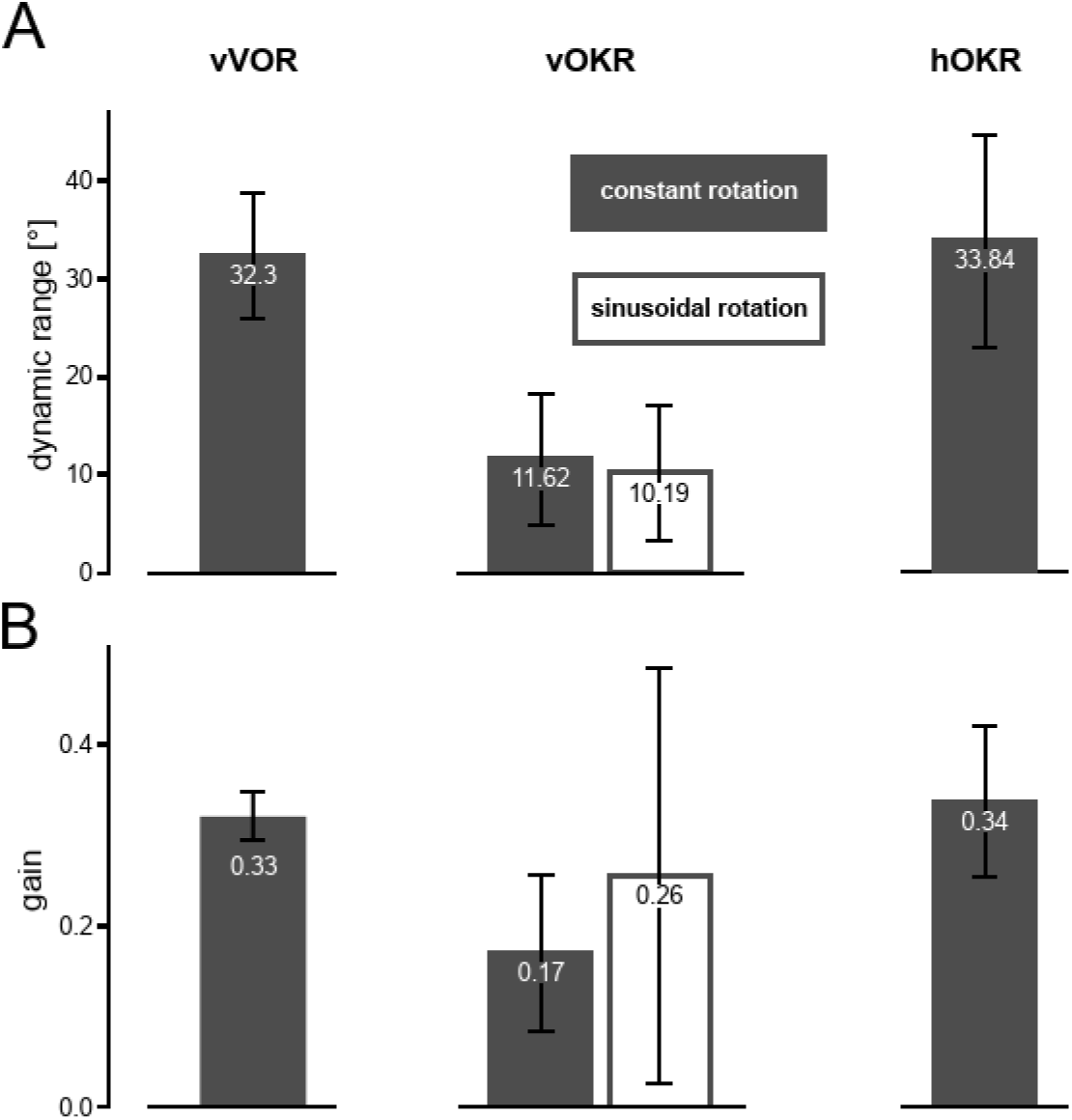
Performance comparison between vertical OKR, vertical VOR, and horizontal OKR. **(A)** Comparison of the dynamic range. For constant rotation (vOKR and hOKR) double the one-directional amplitude was used as dynamic range. For all stimuli, data of recordings with stimulus parameters that elicited the highest gain are shown. **(B)** Comparison of the gain (ratio of eye position to stimulus position after 4 s). For all stimuli, data of recordings with stimulus parameters that elicited the highest gain are shown. All bars show mean values with the standard deviation. For vOKR, filled bars represent values for the constant rotation stimulus whereas open bars represent values for the sinusoidal rotation stimulus.

To better understand the limits and maximal capabilities of the OKR and VOR, we compared the dynamic range and gain across the different reflexes (Figure 5A). While the dynamic range for both constant rotation and sinusoidal vOKR was around 10 °, the dynamic range for vVOR was about three times larger. This striking difference indicates that the vertical deflection range utilized during vOKR is not limited by the maximum possible vertical oculomotor range of the plant and is consistent with recent findings in adult goldfish (Tadokoro et al., 2025). However, the dynamic range of vVOR and hOKR reached similar levels. The gain values for vOKR were smaller compared to hOKR and vVOR, while the gains for hOKR and vVORwere similar (Figure 5B).

## Discussion

This study provides the first characterization of the vOKR in larval zebrafish and compares it to the hOKR and the vVOR. While hOKR has been intensively studied in zebrafish, the vOKR performance remained elusive, even though vertical and torsional OKR have been well-described in various species like primates, cats and even rabbits, which are also lateral-eyed animals, and like zebrafish do not have foveate vision (Collewijn, 1980; Erickson and Barmack, 1980; Farooq, 2004; Kitama et al., 2001; van den Berg and Collewijn, 1988). Since zebrafish serve as a model system for vertebrate visuomotor circuitry, it is essential to know the full behavioral repertoire of the fish. By employing a novel experimental setup, we were able to track both horizontal and vertical eye movements simultaneously while the fish was presented with vertically or horizontally rotating visual stimuli.

### Low vOKR performance levels

In mammals, both the hOKR and vOKR consist of slow phases where the eye follows the stimulus, and resetting quick phases during which the eye rapidly moves in the opposite direction to allow for another slow phase (Erickson and Barmack, 1980; Garbutt et al., 2003; Ghasia and Tychsen, 2014; Robinson, 1981). In our vOKR experiment with a constant rotation stimulus, zebrafish revealed a markedly different response pattern. During the first seconds of stimulation, the fish vertically rotates its eyes in the direction of the stimulus. However, after reaching a certain degree of deflection, the eyes do not move further and do not perform any resetting saccades. This contrasts sharply with a recent study in adult goldfish where regularly occurring resetting saccades during vOKR have been reported (Tadokoro et al., 2025). The authors however mentioned that due to the minimal generated slow phase components, these resetting saccades were unexpected but had excluded any misdetection. This leaves the question whether the absence of any resetting saccades in larval zebrafish vOKR is a species-specific difference or whether they emerge at later developmental stages. Moreover, the zebrafish performs spontaneous saccades at regular intervals during vOKR stimulation, often with a horizontal component. This lack of suppression of spontaneous saccade generation during vOKR is in striking contrast to the vVOR, where not a single saccade occurred during vestibular stimulation. While the vOKR is highly robust in humans, in rabbits it appears to be less effective than the hOKR, with fewer and more irregular resetting saccades, where the eyes often remained at their maximum deflection for over 10 seconds (Erickson and Barmack, 1980).

Generally, the OKR is used to compensate for low-frequency rotations, while the vestibulo-ocular reflex is used for high-frequency rotations of the body (Masseck and Hoffmann, 2009; Robinson, 1981). This principle likely also applies to the vertical domain of the zebrafish sensory-motor systems, although we did not explicitly test it in this study and used different speed regimes for vVOR and vOKR stimulation. We speculate that the vOKR is presumably more relevant for slow and low-frequency roll rotations. Body rotations around this axis probably occur less frequently than for the yaw axis that elicits the hOKR. Such roll rotations are likely quickly adjusted by the gravity-guided posture control mechanisms (Sugioka et al., 2023), which leaves no need for vertical resetting saccades or high dynamic range vOKR responses. Constant visual roll rotation should be very uncommon in healthy fish, which mainly modulate body yaw and - to a lesser extent - body pitch during swimming and navigation. (Budick and O’Malley, 2000; Ehrlich and Schoppik, 2017; Mearns et al., 2020).

Saccades during visual stimulation could be classified into four clusters based on their vertical and horizontal components (Figure S3A and B). Although there are some uncertainties regarding the reliability of the combined horizontal and vertical saccades as large horizontal movements often mask vertical components by projection artifacts in the frontal camera, the presence of pure vertical (Video S3) and pure horizontal (Video S4) saccades alone demonstrates that zebrafish larvae can precisely control their eye muscles to generate saccades with specific vertical and horizontal components. Interestingly, both vertical and horizontal saccades were equally observed during both vOKR and hOKR experiments, although OKR performance was much higher for hOKR (Figure S1B). The vertical saccades during yaw rotation stimuli are likely spontaneous (i.e. not directly driven by the motion stimulus). Further studies are needed to investigate the role of combined vs. pure horizontal and vertical saccades for zebrafish task performance and to reveal the ontogenetic performance levels of older larvae and adult zebrafish. Notably, the transient vOKR during constant rotation displayed a spatial frequency and angular velocity tuning that was highly similar to that previously observed for the hOKR (Dehmelt et al., 2021; Rinner et al., 2005).

The absolute peak gain values of our hOKR differed from a previous study though, with the reported hOKR having about three-times higher gain levels than our hOKR at similar angular velocities and spatial frequencies (Rinner et al., 2005), which can potentially be explained by the smaller steradian size of our stimulus arena. Nevertheless, the tuning curve shape of hOKR to velocity and spatial frequency is similar. The dynamic range was significantly different between the vertical and horizontal OKR, with the hOKR reaching much larger eye positions. This is particularly interesting given that during the vVOR, zebrafish were able to achieve three-times absolute higher vertical eye positions than during vOKR. Together with the high initial vOKR gain and the reported vertical motion sensitivity in the pretectum (Wang et al., 2019), these findings suggest that the limiting factor for vOKR is neither visual motion sensitivity nor the oculomotor plant. Instead, these results point towards the inability of diencephalic direction-selective neurons in recruiting high levels of motoneuron pool activity needed for very peripheral eye positions. This could be caused by motoneurons with high eye position thresholds only being recruited by vestibular and not by visual input (Aksay et al., 2000; Brysch et al., 2019) or by the presence of saccade-type specific motoneuron populations (Brysch et al., 2019; Dowell et al., 2025; Leyden et al., 2021).

### Phase advances during low repetition rates might be related to adaptation or acceleration tuning

Using the sinusoidal rotation stimulus, it was possible to evoke a long-lasting sinusoidal vOKR that followed the stimulus position. For low repetition rates that resulted in extended periods of unidirectional stimulus motion, the eyes reached their maximum vertical deflection already after several seconds and remained at that plateau. One might expect the eyes to stay at the plateau until the stimulus changed direction, at which point the eyes would move in the opposite direction away from the plateau. However, the eyes appeared to start rotating in the opposite direction several seconds before the stimulus changed direction, i.e. the stimulus was still moving in the original direction (just slower). This was true for all repetitions during a stimulation phase and did not evolve over time (Figure S2C), which is why predictive behavior occurring after a training period with a repetitive stimulus can be ruled out (Marsh and Baker, 1997; Miki et al., 2018).

This phenomenon shares some similarities with the motion aftereffect (MAE), an illusion of motion in a certain direction caused by prior exposure to motion in the opposite direction (Anstis et al., 1998). The motion aftereffect can reliably be elicited by moving visual stimuli (Braun et al., 2006; Chen et al., 2014; Watamaniuk and Heinen, 2007) and is also known to be present in larval zebrafish with regard to both the optomotor and optokinetic response (Lin et al., 2019; Najafian et al., 2014; Pérez-Schuster et al., 2016; Wu et al., 2020). After prolonged hOKR stimulation, fish exhibit a hOKR in the opposite direction during a stationary stimulus, and the duration of the reversed response depends on the duration of the initial stimulation. At least 200 seconds of hOKR stimulation is necessary to induce an MAE according to Pérez-Schuster et al. (Pérez-Schuster et al., 2016). Two key differences to the behavior observed in this study are that i) zebrafish began moving their eyes in the opposite direction while the stimulus was still moving, and ii) that our observed phase-advanced vOKR occurred for stimulation periods much shorter than 200 seconds. For the lowest repetition rate, the duration of stimulus rotation in one direction was over 2 minutes, whereas for a repetition rate of 0.125 Hz, the stimulus rotated for only 4 seconds in one direction, and a small phase advance was already visible. One potential explanation for the occurrence of the phase advance could be an interplay between two mutually inhibiting populations of direction-selective (DS) neurons coding for opposite velocities. The population coding for the stimulus direction may adapt over time, until the difference between the two population’s activities is so small that a deceleration of the stimulus disinhibits the non-stimulated DS population, which induces a vOKR in the opposite direction. However, the fast adaptation (< 4 s for 0.125 Hz repetition rate) is much faster than previously reports on MAE in zebrafish (Pérez-Schuster et al., 2016; Wu et al., 2020).

An alternative explanation for the occurrence of phase advances is that the DS neurons - next to their tuning to velocity - are tuned to acceleration as well (Cao et al., 2004; Thiel et al., 2007), which can be interpreted as a quick form of adaptation, and thus they might lose much of their activity when the stimulus is still running in the same direction but acceleration went missing. In this scenario, the occurring eye movements into the opposite direction (back to the null position) would not necessarily be visually driven, but could simply result from passive drift back of the eyes in the oculomotor plant (Miri et al., 2022). In line with this hypothesis, centripetal eye movements (which can be interpreted more safely as visually driven) only occurred after the stimulus velocity had switched sign (Figure 3D), i.e. vOKR towards peripheral positions was always going into the same direction as the stimulus.

Further studies are needed to determine the mechanisms that underlie the observed apparent phase advance of vertical optokinetic responses to slowly repeating sinusoidal rotation stimuli.

### Uncertainties associated with projection-based eye tracking

The dual-camera setup used for all recordings enabled simultaneous tracking of horizontal and vertical eye movements in zebrafish larvae. However, many eye movements were visible in both horizontal and vertical eye traces. In particular saccades with large amplitudes were often visible in both cameras. We tracked horizontal and vertical eye movements in larval zebrafish by fitting an ellipse to the binarized eye image and calculating the angle between the major axis and the body midline. This method should be relatively robust as compared to a previously reported method based on searching the largest distance of two pixels within a binarized and segmented eye (Dehmelt et al., 2018) and has also been used to measure vertical eye positions in a previous study (Bianco et al., 2012).

This method can reliably track both pure vertical and pure horizontal eye movements, which can be confirmed by inspecting raw eye traces and measurements such as the change in length of the eye’s fitted minor axis during saccadic eye movements and difference images during saccades (see above). However, the method raises some uncertainties regarding combined vertical and horizontal eye movements. While a large amount of eye movements are visible in both the horizontal and vertical planes, the method reaches its limits when attempting to disentangle the vertical and horizontal components of apparently mixed eye movements. Large horizontal eye movements often (but not always) lead to a mis-detection of vertical eye position in the frontal camera, caused by an ill-defined ellipse fit with a major/minor axis ratio closer to one than for eye positions without horizontal components. This change in projection shape could affect the calculated vertical angle and therefore mask the true vertical component of the movements. The change in projection shape in the frontal camera was especially apparent during hOKR recordings, where the stereotypical slow/quick phase pattern was also observed in the vertical eye trace, albeit with a much smaller amplitude. This even resulted in a small apparent tuning of vertical eye movements to hOKR stimulation (Figure S1B). If these eye movements were genuine, it would suggest that the hOKR had a constant vertical component, which would be surprising, even considering the potential for small body pitch due to imprecise embedding of the larva. Nevertheless, combined vertical and horizontal eye movements are found to be present in adult goldfish (Tadokoro et al., 2025) and are most likely also present in zebrafish.

Taken together, our data shows that pure horizontal and pure vertical saccades do exist, as well as combined movements, although the true vertical and horizontal components of possible mixed saccades could not be revealed in this study. The precise measurement of vertical components in apparent horizontal-vertical saccades is hampered by methodological caveats, and a different tracking method of the dual-camera recordings needs to be performed in the future to improve our understanding of the horizontal and vertical components of mixed-saccades in larval zebrafish.

## Conclusion

In this study, we characterized the vOKR in larval zebrafish and compared its characteristics to the hOKR and the vVOR. Larval zebrafish do perform vOKR and several basic differences from the hOKR could be identified. One major difference was the absence of resetting vertical saccades during vOKR. The reason for this needs to be explored in future studies. Given that vertical rotation encoding neurons exist (Wang et al., 2019; Zhang et al., 2022), a better understanding of the downstream neuronal processes controlling eye movement and saccade generation could help explain the behavioral differences between vertical and horizontal OKR. Such neurophysiology experiments could also shed light on the differences in dynamic range between vOKR and the vVOR. Additionally, further behavioral experiments are necessary to understand the apparent vOKR phase advance for sinusoidal rotation stimuli in the context of adaptation or acceleration tuning. Our characterization of the vOKR in larval zebrafish provides information on a hitherto elusive behavior and points towards specific questions that will help gain a more comprehensive understanding of the neuronal processing of visual information and the control of eye movements.

## Supporting information

supplementary KEBAB

## Author Contributions

D.-S.B. and T.C.H. developed and built the setups with the help of C.F. D.-S.B., G.M. and A.B.A. developed the methodologies. G.M., L.D. and C.F. recorded all data. G.M. and D.- S.B. did the data analysis and made the figures. D.-S.B., A.B.A and G.M. wrote the manuscript. D.-S.B. and A.B.A. supervised the project. A.B.A. provided all resources.

## Data Availability

All data are available on request.

