## supplementary figures for "Vertical Optokinetic Eye Movements in the Larval Zebrafish"

### Supplementary Figures and Tables

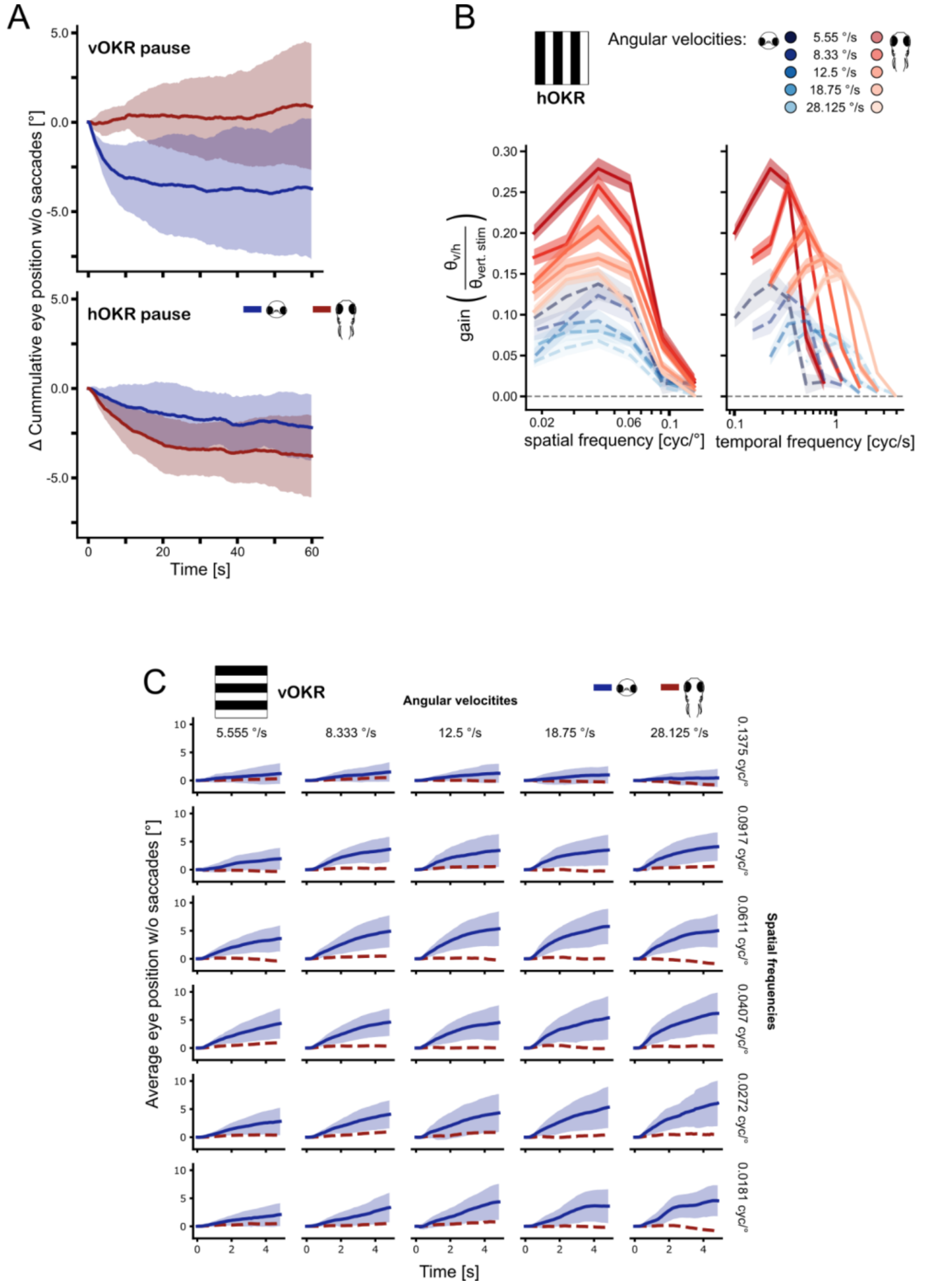


**Figure S1: Additional data for constant rotation OKR stimulus analysis. (A)** Relative Average cumulative eye position with saccades removed for pause breaks during vOKR (top) and hOKR (bottom) stimulus protocols. Depending on the rotation direction of the previous stimulation, the individual eye traces have been inverted. **(B)** Gain tuning to spatial frequency and temporal frequency for all different angular velocities. Lines show mean and standard error of the mean. Red: gain for horizontal eye trace for hOKR stimulation. Blue: gain calculated for vertical eye trace for hOKR stimulation. Dashed horizontal line indicates a gain of 0. **(C)** Average start response during the first seconds of vOKR stimulation for all stimulus combinations. Blue line indicates average vertical eye trace and envelope indicates standard deviation. Red dashed line indicates the average horizontal eye trace. n = 10.

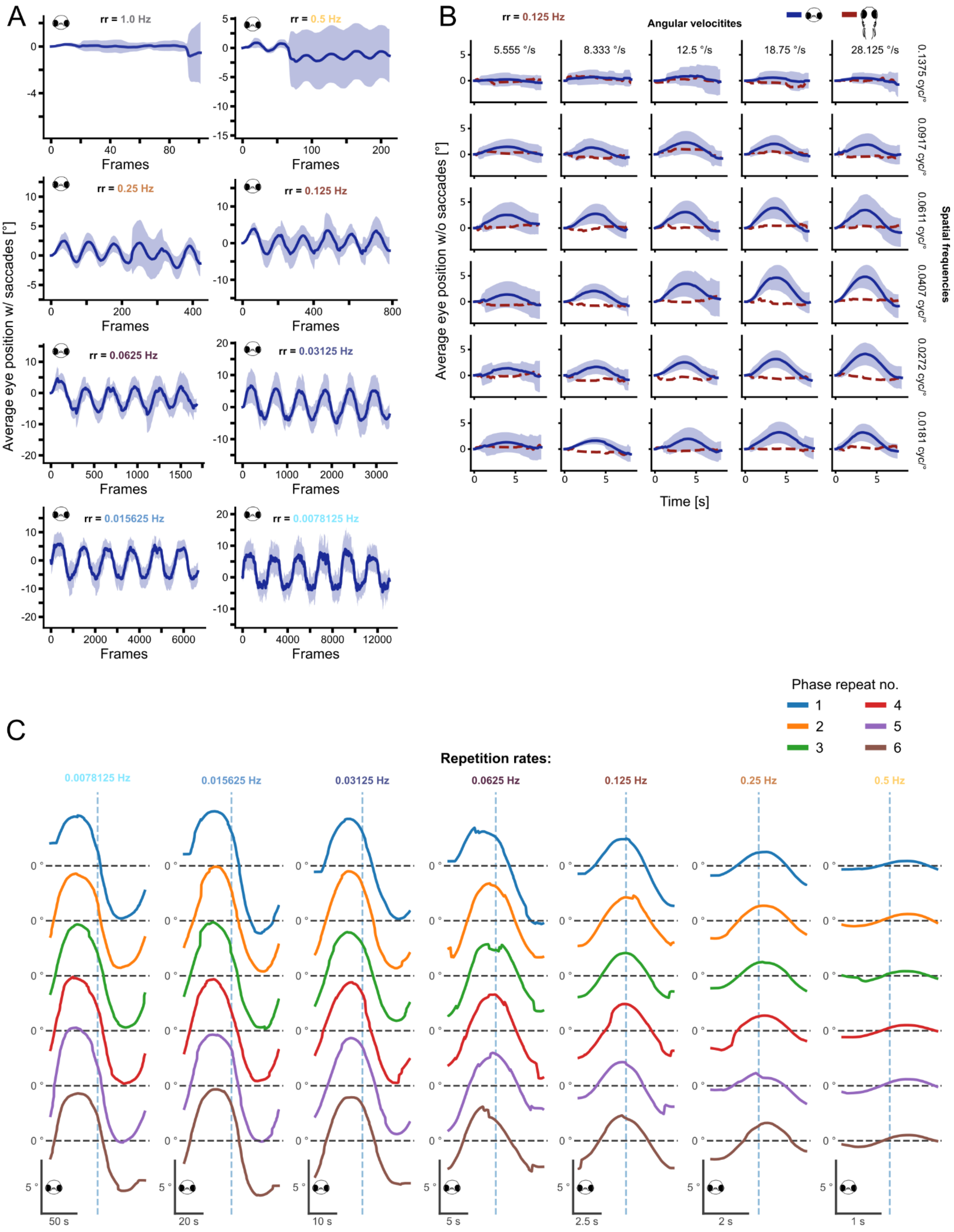


**Figure S2: Additional Data for sinusoidal vOKR. (A)** Averaged eye positions for the entire stimulation with each repetition rate (rr) in the rr-tuning sinusoidal vOKR stimulus without saccade removal. First responses are elicited by rr = 0.5 Hz. For rr = 0.0625 Hz and lower, the eyes reach a maximum deflection which becomes more distinct for lower rr. Increase in standard deviation for 1.0, 0.5, and 0.25 Hz is due to individual fish that performed a large saccade during the stimulation and thereby shifted the eye trace into one direction. n = 8. **(B)** Average response to the stimulus (stimulus-triggered average, STA) for all stimulus combinations of the fs-tuning sinusoidal vOKR stimulus. Blue line indicates vertical STA and envelope indicates standard deviation. Red dashed line indicates the horizontal STA. n = 11. **(C)** Individual average responses to each stimulus repetition for the different repetition rates of the rr-tuning vOKR stimulus. Dashed vertical line indicates midpoint of one stimulation phase as reference. Phase lag is consistent over all repeats within the same stimulus condition. n = 8.

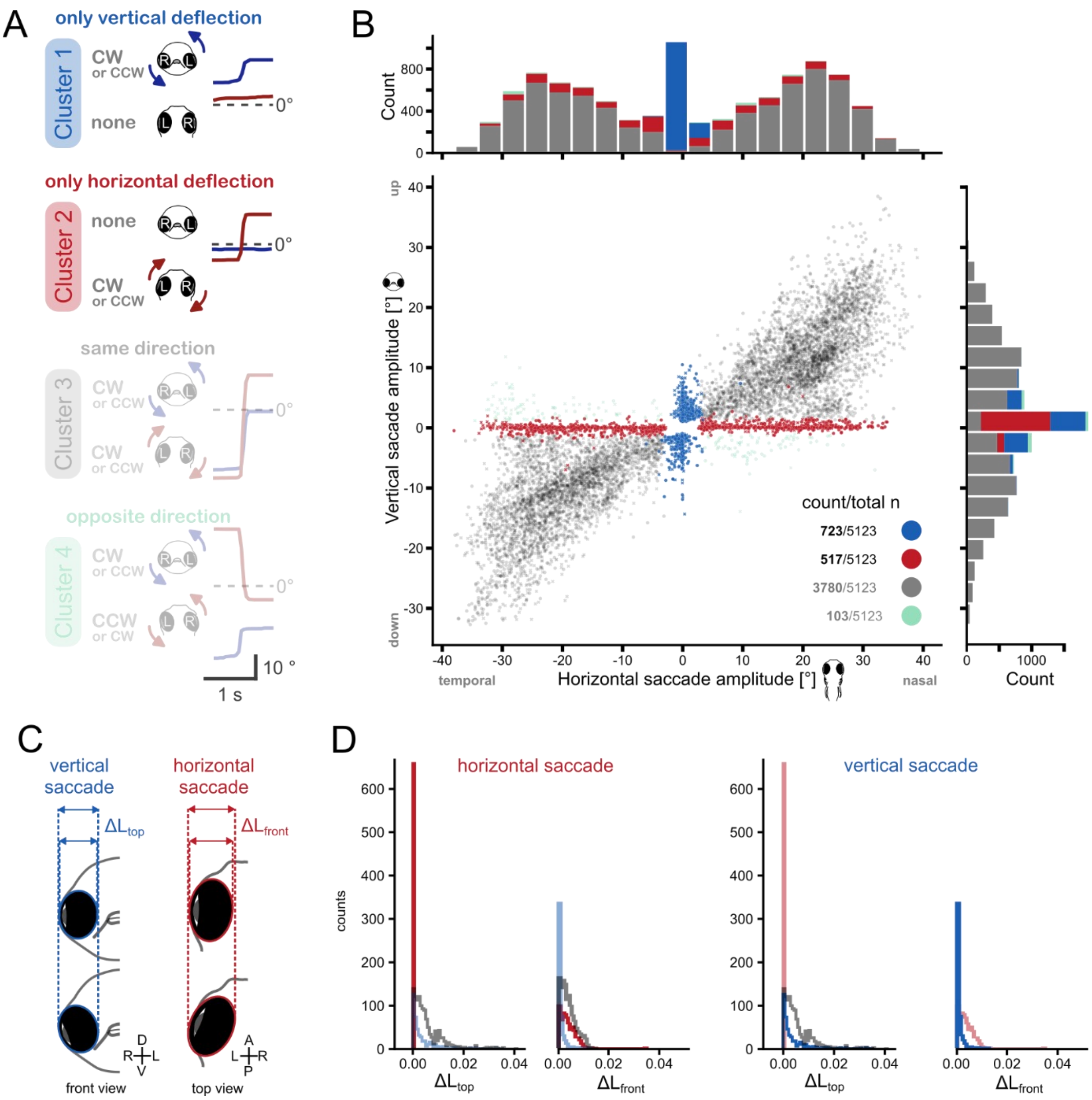


**Figure S3: Saccade Clusters during vertical OKR (vOKR) stimulation. (A)** Four different saccade types have been observed and can be distinguished by their appearance in the vertical and horizontal eye trace. Example vertical (blue) and horizontal (red) eye traces show characteristics of each cluster type. Scale on the right corresponds to all eye traces. Note, that cluster 3 and 4 contain both mixed (vertical-horizontal) and pure horizontal saccades. Pure horizontal saccades can cause projection artifacts on the front camera (due to large movement, illumination and eye position) which can result in an apparent vertical movement. Cluster 3 and 4 were therefore not further analyzed.

**(B)** Scatterplot of apparent horizontal and vertical amplitudes of all observed saccades, clustered by the saccade type during vOKR stimulus. **(C)** Illustration of the minor axis length change (after ellipse fitting) during pure vertical and horizontal saccades. **(D)** Distribution of minor axis length changes during horizontal (red) and vertical (blue) saccades for both top ($\Delta L_{front}$) and front ($\Delta L_{top}$) view. Red and blue bars represent the minor axis length change distributions during horizontal and vertical saccades. Grey bars represent distributions during mixed saccades. During a horizontal saccade, the minor axis length changes of the top view are ~0 whereas the minor axis length changes of the front view are >0 (left). For vertical saccade, it is exactly the opposite (right). This measurement confirms detection of true horizontal and vertical saccades, as the minor axis length of the top view ($L_{front}$) should not change during a horizontal saccade, but should change during a vertical saccade and vice versa for the minor axis length of the front view ($L_{top}$).

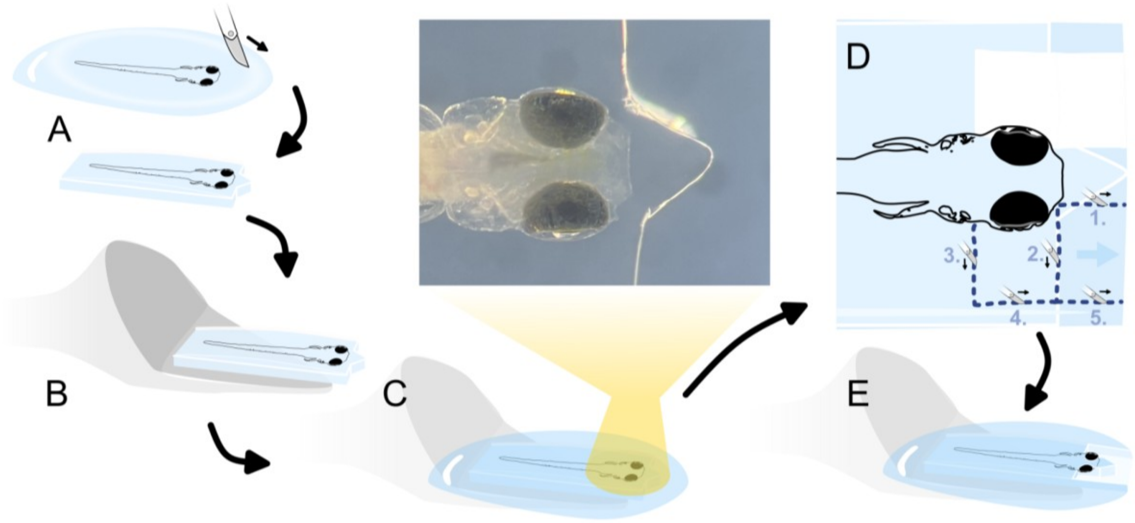


**Figure S4: Schematic of zebrafish larva embedding procedure. (A)** The larva is immobilized in 1.6 % agarose and a block containing the fish is cut out. The image shows the triangle shape of the first agarose block in front of the head of the fish. **(B)** The block containing the fish is transferred onto a modified pipette tip. **(C)** The agarose block with the zebrafish is fixated on the pipette tip with agarose. **(D)** Scheme of steps to remove the agarose around the eyes. **(E)** The fish is ready for recording; the eyes can move and have no agarose in front of them while a small block of agarose remains in front of the head to prevent the fish from freeing itself.

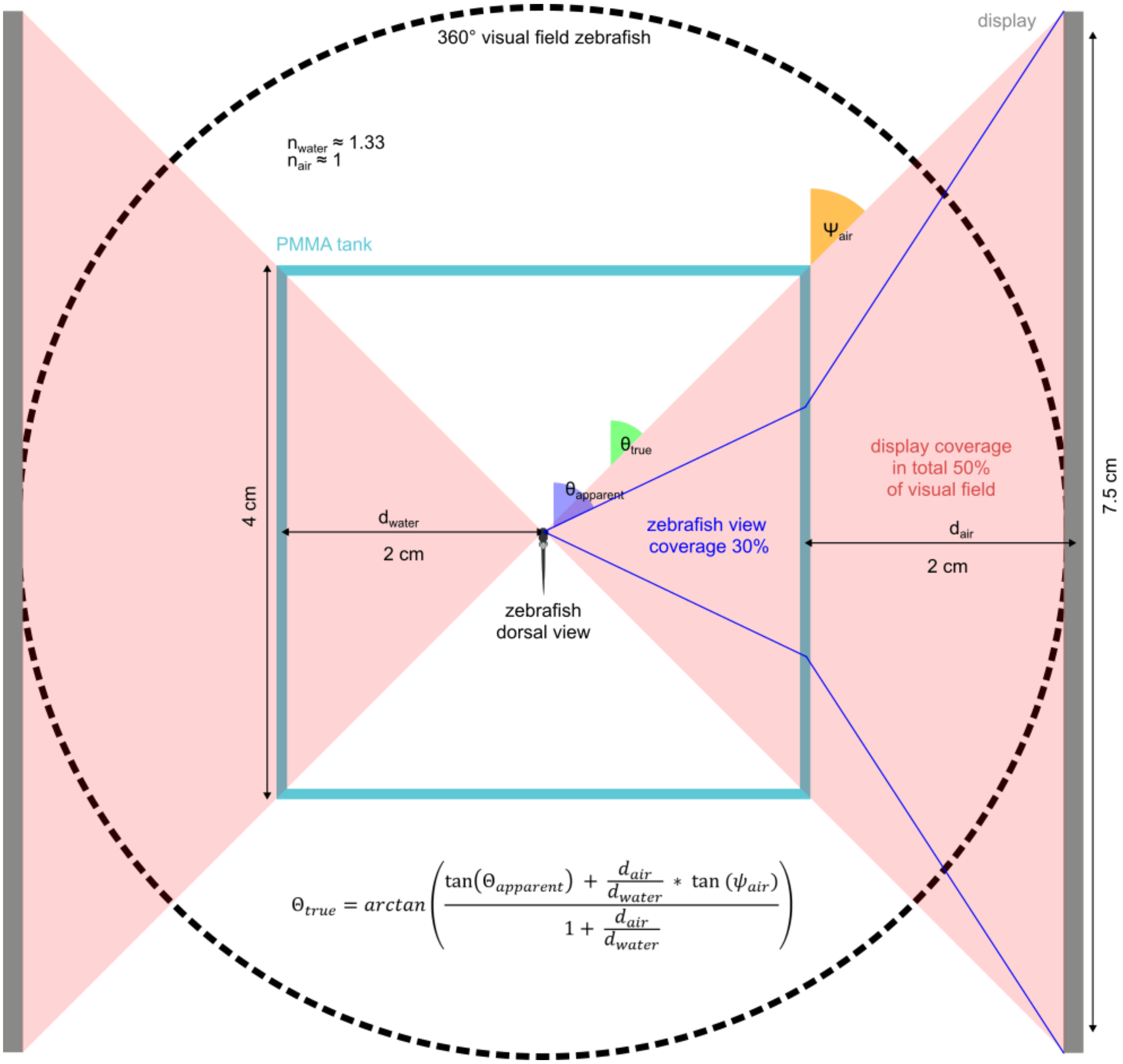


**Figure S5: Dimensions and visual coverage of OKR setup. (A)** Dimension of the visual stimulation setup. Visual coverage of the arena was calculated according to Dunn and Fitzgerald (2020). Red area marks the visual coverage of the displays without correcting for physical distortions. Blue lines mark the area of visual coverage after correcting for refractive distortions.

| **Spatial Frequency [cycle/°)** | **Spatial Period [°/cycle]** | **Angular Velocity [°/s]** | **Temporal Frequency [cycle/s]** |
|---|---|---|---|
| 0.0181 | 55.2486 | 28.125 | 0.77 |
| | | 18.75 | 0.51 |
| | | 12.5 | 0.34 |
| | | 8.333 | 0.23 |
| | | 5.555 | 0.15 |
| 0.0272 | 36.7647 | 28.125 | 0.77 |
| | | 18.75 | 0.51 |
| | | 12.5 | 0.34 |
| | | 8.333 | 0.23 |
| | | 5.555 | 0.15 |
| 0.0407 | 24.5700 | 28.125 | 1.14 |
| | | 18.75 | 0.76 |
| | | 12.5 | 0.51 |
| | | 8.333 | 0.34 |
| | | 5.555 | 0.23 |
| 0.0611 | 16.3666 | 28.125 | 1.72 |
| | | 18.75 | 1.15 |
| | | 12.5 | 0.76 |
| | | 8.333 | 0.51 |
| | | 5.555 | 0.34 |
| 0.0917 | 10.9051 | 28.125 | 2.58 |
| | | 18.75 | 1.72 |
| | | 12.5 | 1.15 |
| | | 8.333 | 0.76 |
| | | 5.555 | 0.51 |
| 0.1375 | 7.2727 | 28.125 | 3.87 |
| | | 18.75 | 2.58 |
| | | 12.5 | 1.72 |

| | | | |
|---|---|---|---|
| | | 8.333 | 1.15 |
| | | 5.555 | 0.76 |

**Table S1: Stimulus parameters for assessing OKR tuning to spatial frequency and angular velocity.** Each stimulus parameter pair was repeated for both rotation directions. Parameter values were adapted from (Dehmelt et al., 2021). The order of the stimulus parameter pairs was randomized.

| Spatial frequency [cycle/°] | Spatial period [°/cycle] | Angular velocity [°/s] | Repetition rate [cycle/s] | Length for 6 sine period repetitions [s] |
|---|---|---|---|---|
| 0.0611 | 16.3666 | 12.5 | 3.0 | 2 |
| | | | 2.0 | 3 |
| | | | 1.0 | 6 |
| | | | 0.5 | 12 |
| | | | 0.25 | 24 |
| | | | 0.125 | 48 |
| | | | 0.0625 | 96 |
| | | | 0.03125 | 192 |
| | | | 0.015625 | 384 |
| | | | 0.0078125 | 768 |

**Table S2: Stimulus parameters for assessing vOKR repetition rate tuning.** The stimulus was presented two times for each repetition rate and with a randomized direction of the maximum angular velocity.

**Video S1: Example recording of vVOR experiment.** Recording of vestibular stimulation protocol using the KEBAB setup with a velocity of 90°/s and constant rotation. The compensatory vertical eye movements are clearly visible in the camera (left side of video) and the corresponding eye traces of live tracking (using ellipse fitting on thresholded eye) display a sinusoidal pattern, visible in the right side. To ensure maximal eye tracking performance, image parameters like contrast and brightness were adjusted.

**Video S2: Example recording of sinusoidal vOKR experiment.** Recording of sinusoidal visual stimulation using the two-camera visual stimulation setup. In the top half, both camera views (top and front) are visible. To ensure a good eye tracking performance, image parameters like contrast and brightness were adjusted. Note: The front image (right top corner) was rotated 90° due to the camera position. Pure vertical sinusoidal eye movements (vOKR) are clearly visible in both top and front camera view. The lower half shows the corresponding eye traces of live tracking. Bright and dark blue lines correspond to vertical eye movements and bright and dark orange lines correspond to horizontal eye movements. Note: The video was sped up to allow visual inspection of different phases with different repetition rates (as can be seen by the different frequencies of the sinusoidal vOKR).

**Video S3: Pure vertical saccade.** Video of front (left) and top (right) view during a pure vertical saccade.

**Video S4: Pure horizontal saccade.** Video of front (left) and top (right) view during a pure horizontal saccade.
