## supplementary KEBAB for "Vertical Optokinetic Eye Movements in the Larval Zebrafish"

### KEBAB Construction Overview

#### **K**inematic **E**valuation of **B**ehavior during **A**nimal **B**ody-roll

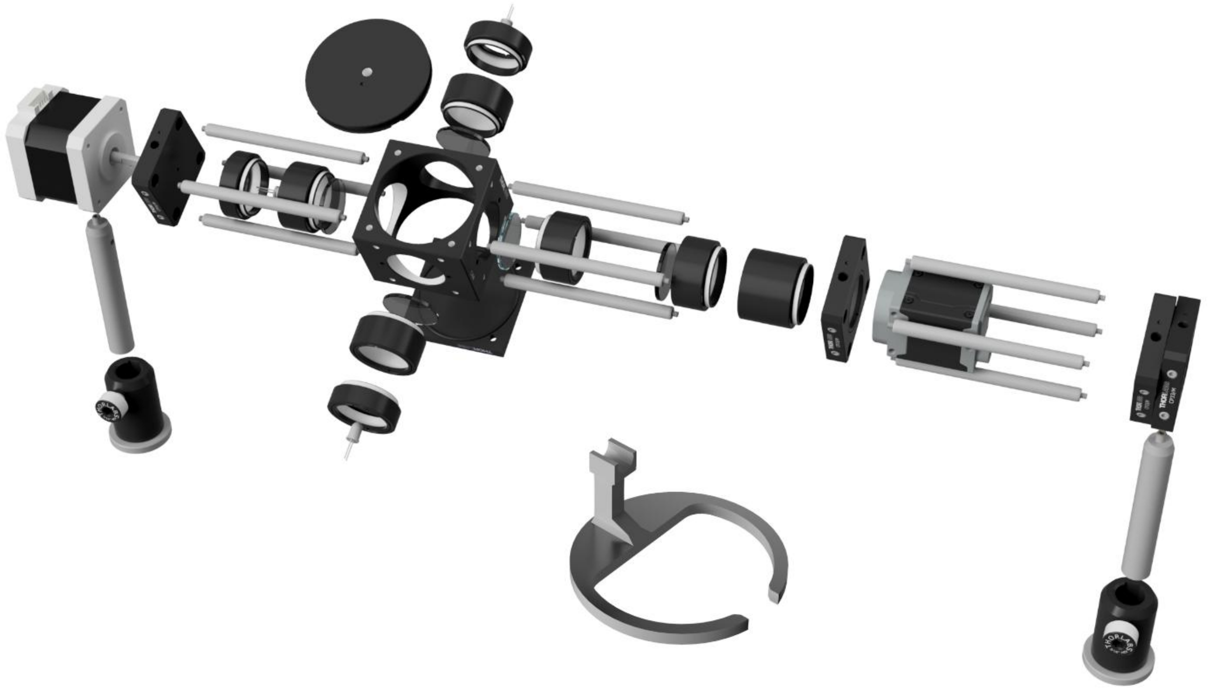

**Wiring Layout:**

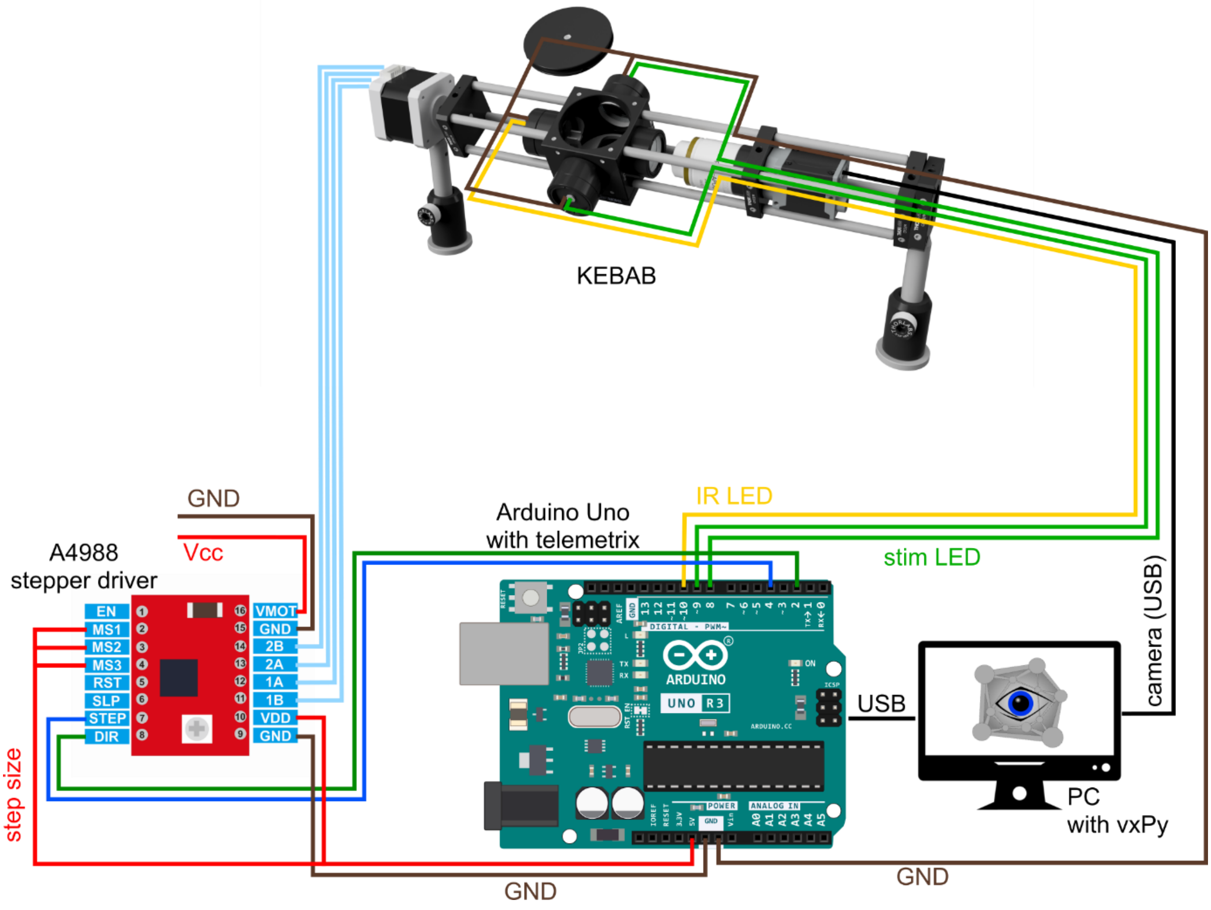

**Software:**

vxPy: https://github.com/thladnik/vxPy

telemetrix: https://mryslab.github.io/telemetrix/

**Components:**

1x NEMA 17 stepper motor, ACT Motor GmbH

5x SM1 Lens Tubes 0.5”, Thorlabs

3x SM1 Lens Tubes 0.3”, Thorlabs

3x 1” UV Fused Silica Ground Glass Diffusor, Thorlabs

1x 1” UVFS Broadband Precision Window, Thorlabs

1x 30mm Rotating Cage Segment Plate, Thorlabs

1x Blank 30mm Cage Plate with bore for NEMA 17 axle, Thorlabs

1x Blank Cover Plate, Thorlabs

1x 30 mm Cage Cube, Thorlabs

1x Fixed Cage Cube Platform for C4W/C6WR, Thorlabs

1x SM1-Threaded 30mm Cage Plate, Thorlabs

12x 3” Cage Assembly Rods for 30mm Cage, Thorlabs

2x 0.5” Optical Post, Thorlabs

2x 0.5” Pedestal Post Holder, Thorlabs

1x 0.48” Diameter 12 Wire Capsule Slip Ring, Amazon

1x IR LED 850 nm, e.g. Conrad

2x white LED, e.g. Conrad

1x 1” IR 850nm longpass filter, Thorlabs

1x camera DMK 23UV024, Imaging Source

1x C-mount 5x extender lense, e.g. bestscientific

1x Arduino Uno, Conrad

1x A4988 stepper driver, Conrad

Noise floor measurements:

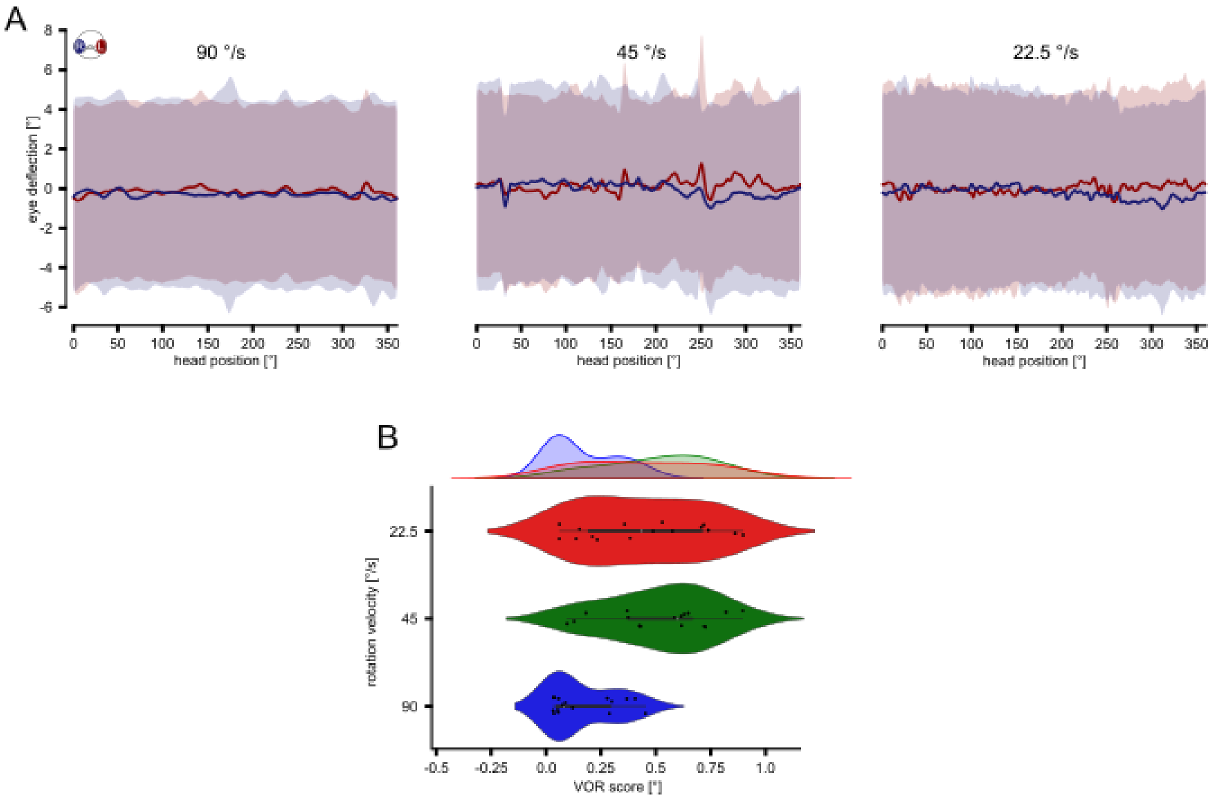


Noise measurements. Measurements of anesthetized 4dpf larvae (168mg/l MS222) running the same VOR protocol as used in our main study. A) STA of eye movements during different rotation velocities (90 °/s, 45 °/s, 22.5 °/s). B) VOR score of anesthetized larvae for different rotation velocities (90 °/s, 45 °/s, 22.5 °/s).
